# Advancing Pediatric and Longitudinal DNA Methylation Studies with CellsPickMe, an Integrated Blood Cell Deconvolution Method

**DOI:** 10.1101/2025.04.22.649907

**Authors:** Maggie P. Fu, Karlie Edwards, Erick I. Navarro-Delgado, Sarah M. Merrill, Negusse T. Kitaba, Chaini Konwar, Piush Mandhane, Elinor Simons, Padmaja Subbarao, Theo J. Moraes, John W. Holloway, Stuart E. Turvey, Michael S. Kobor

**Affiliations:** Edwin S. H. Leong Centre for Healthy Aging, Faculty of Medicine, University of British Columbia, Vancouver, BC, Canada; Centre for Molecular Medicine and Therapeutics and Department of Medical Genetics, University of British Columbia, Vancouver, BC, Canada; British Columbia Children’s Hospital Research Institute, Vancouver, BC, Canada; Department of Psychology, University of Massachusetts Lowell, Lowell, MA, USA; Human Development and Health, Faculty of Medicine, University of Southampton, Southampton, UK; Department of Pediatrics, University of Alberta, Edmonton, AB, Canada; Department of Pediatrics & Child Health, University of Manitoba, Winnipeg, MB, Canada; Translational Medicine Program, The Hospital for Sick Children, Toronto, ON, Canada; NIHR Southampton Biomedical Research Centre, University Hospitals Southampton, Southampton, UK; Department of Pediatrics, The University of British Columbia, Vancouver, Canada

## Abstract

Prospective birth cohorts offer the potential to interrogate the relation between early life environment and embedded biological processes such as DNA methylation (DNAme). These association studies are frequently conducted in the context of blood, a heterogeneous tissue composed of diverse cell types. Accounting for this cellular heterogeneity across samples is essential, as it is a main contributor to inter-individual DNAme variation. Integrated blood cell deconvolution of pediatric and longitudinal birth cohorts poses a major challenge, as existing methods fail to account for the distinct cell population shift between birth and adolescence. In this paper, we critically evaluated the reference-based deconvolution procedure and optimized its prediction accuracy for longitudinal birth cohorts using DNAme data from the Canadian Healthy Infant Longitudinal Development (CHILD) cohort. The optimized algorithm, *CellsPickMe*, integrates cord and adult references and *picks* DNA*me* features for each population of *cells* with machine learning algorithms. It demonstrated improved deconvolution accuracy in cord, pediatric, and adult blood samples compared to existing benchmark methods. *CellsPickMe* supports blood cell deconvolution across early developmental periods under a single framework, enabling cross-time-point integration of longitudinal DNAme studies. Given the increased resolution of cell populations predicted by *CellsPickMe*, this R package empowers researchers to explore immune system dynamics using DNAme data in population studies across the life course.

## Introduction

The Developmental Origins of Health and Disease (DOHaD) hypothesis posits that exposures during sensitive developmental windows, most especially within the first years of life, can influence long-term physiological functioning and the health trajectory of a person^1,2^. DNA methylation (DNAme), an epigenetic mark that is mitotically heritable but malleable to environmental changes, has been proposed as a molecular process that contributes to the biological embedding of early life experiences^3–8^. Pregnancy and birth cohorts have been established with longitudinal follow-up with deep phenotyping, including DNAme profiling, to examine the association between environmental exposures and health outcomes later in life^9–13^.

However, this early developmental period coincides with profound remodeling of the hematopoietic landscape^14–16^. This includes a rapid loss of nucleated red blood cells (nRBCs) after birth^17–19^, increased neutrophil proportion accompanied with high neutrophil heterogeneity in neonates^20–22^, and changes in naïve and memory lymphocyte profiles upon antigen exposure including vaccinations^14,15,23,24^. The dynamic of peripheral blood cell populations provides insight into individuals’ immune status, stress response, as well as developmental and aging trajectory^25–28^; yet, it creates an additional layer of complexity to epigenomic investigations, as cellular functions and identities are intrinsically linked to their DNAme signature^29–31^

Longitudinal birth studies have reported persistent and wide-spread alterations in DNAme within the first years of life^32–34^. These studies compared the DNAme patterns of cord blood with that of peripheral blood collected at pediatric time points, ranging from neonates (less than 1 month old), infants (birth – 2 years), children (2 – 12 years), to adolescents (12 – 21 years)^32–35^. In this context, cellular composition of a heterogeneous tissue like blood can be bioinformatically inferred without readily available cell count data, using DNAme-based deconvolution methods^36^. Several cell type deconvolution algorithms have been independently developed for adult and cord blood samples using a reference-based approach, leveraging cell-type specific DNAme signatures from sorted cells^36–40^. However, without a method to harmonize deconvolution of cord, pediatric, and even adult blood under a single framework, direct inference of DNAme changes across the early developmental period is inevitably confounded by a profound variance in cellular composition^41–44^.

To address this methodological gap, it is crucial to note that not only does the cellular composition of blood varies with age, but so does the DNAme profile of sorted cells^37,45–47^. The findings suggest that applying a reference dataset developed from cord blood or adult blood in pediatric populations may introduce prediction errors^48^. Additionally, performing cellular deconvolution in longitudinal birth cohorts spanning a range of ages currently require multiple, non-comparable references. To overcome this conundrum, we proposed an integrated DNAme reference of cord and adult sorted blood cells, with the goal of capturing cell-type-specific signals that account for changes in the hematopoietic landscape. In addition to reference dataset selection, we evaluated the impact of other steps in the reference-based cell type deconvolution procedure, including data normalization, feature selection, and regression-based prediction^49^. The algorithm’s performance was assessed using the pediatric blood samples of a Canadian population-based prospective birth cohort, the Canadian Healthy Infant Longitudinal Development (CHILD) cohort^11^. The resulting reference-based deconvolution procedure, *CellsPickMe*, utilizes machine learning-based method to *pick* DNA*me* features for each *cell population*. *CellsPickMe* with the curated UniBlood references facilitates the joint deconvolution of cord and whole blood and supports the inference of cell type proportions in longitudinal pediatric cohorts where age-appropriate reference datasets are currently unavailable.

## Results

### Cellular maturity and developmental time point covaries with DNAme pattern

We first explored whether there is a distinct DNAme signature across developmental stages that can impact pediatric cellular deconvolution process. To examine DNAme variation through the life course, we applied principal component analysis (PCA) to sorted DNAme reference data sets of cord and adult blood: *FlowSorted.CordBlood.450k* (referred as Cord), *FlowSorted.Blood.EPIC* (referred as IDOL), and *FlowSorted.Blood.Extended.EPIC* (referred as Extended). Immune cells clustered distinctly on both a lineage axis along PC1(myeloid to lymphoid) and a maturity axis along PC2 (neonatal – naïve – memory) (**Figure 1A**). The nRBCs showed a distinct DNAme pattern, clustering in the middle of lymphocytes and granulocytes on the lineage axis, and further down the maturity axis compared to other cord immune cells. For T and B cells, there was an evident gradient along the maturity axis, transitioning from neonatal, adult naive, a mixture of adult naive and memory (IDOL reference did not distinguish between the two), and adult memory cells (**Figure 1A**).

**Figure 1.**
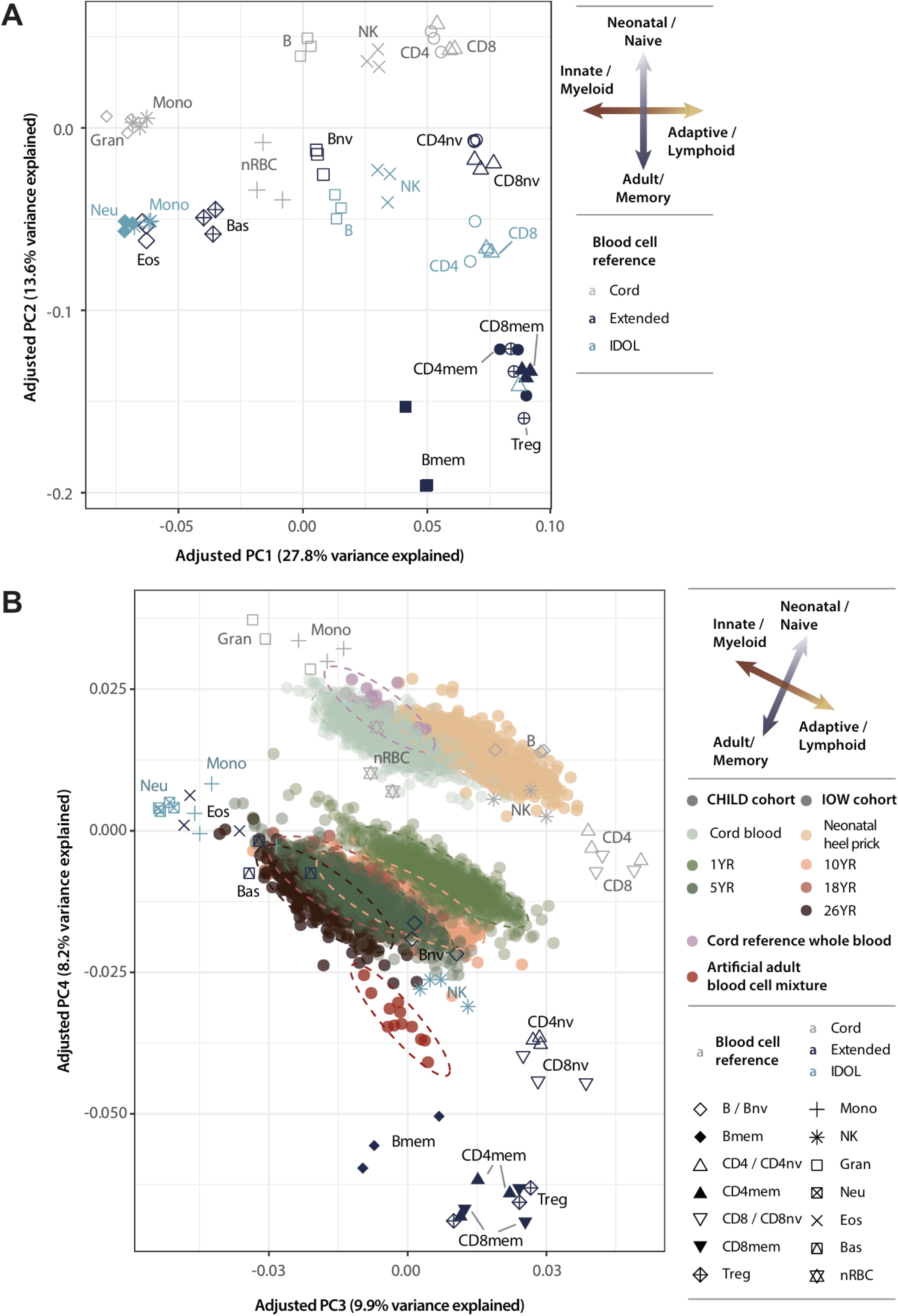
Principal component analysis showing the main patterns of DNAme variation in **A)** sorted cord and adult peripheral blood cell populations (plotted as text indicating their cell type) and **B)** the sorted cells from **A** with umbilical cord blood from Cord reference (pink dots), artificial mixture of adult blood cells (red dots), and two additional longitudinal pediatric cohorts: CHILD (green dots) and IOW (orange-brown dots). The ellipses were created based on 95% confidence interval using the *ggplot2:: stat_ellipse()* function. In both **A** and **B**, three of each sorted cell type were randomly sampled and shown to reduce visual cluster. nv: Naïve cells; mem: Memory cells; Treg: Regulatory T cells; NK: Natural killer cells; Mono: Monocytes; Gran: Granulocytes; Neu: Neutrophils; Bas: Basophils; Eos: Eosinophils; nRBC: Nucleated red blood cells.

To study whether the pattern of DNAme variation is recapitulated in whole blood where bulk DNAme is representative of all cell types present in the sample, we clustered the DNAme profiles of sorted immune cells with that of cord and whole blood from two longitudinal birth cohorts, the Isle of Wight (IOW) and the Canadian Healthy Infant Longitudinal Development (CHILD) cohorts, along with an artificial blood cell mixture from adult sorted cells, and whole umbilical cord blood from the Cord reference. Aside from technical batch effect and biological sex, which contributed to PC1 and PC2 variance, we observed an analogous trend of lineage and maturity axes along PC3 and PC4 (**Figure 1B**). The heterogeneous blood samples clustered in the middle of PC3, as they are composed of a mixture of myeloid and lymphoid cells. These included the CHILD and IOW’s post-natal blood, the artificial blood mixture and the Cord reference cord blood samples. The CHILD cord blood showed a high overlap with the CORD samples and nRBC cells, whereas the IOW neonatal dried blood spots, with a reduced nRBC fraction, clustered adjacently as previously observed^50^. As the CHILD and IOW participants aged, their samples shifted down the maturity axis (**Figure 1B**). At age 1, the CHILD samples clustered discretely from both cord and adult samples. As individuals aged, age 5 CHILD samples and age 10 and 18 IOW samples became more similar, yet remained in a distinct cluster from the age 26 IOW and ABM samples.

### Optimization procedure of reference-based cell type deconvolution

Building on evidence indicating that developmental stages in childhood are linked to distinct DNAme profiles, we aimed to refine the cell type deconvolution process for pediatric blood samples. **Figure 2** shows a flowchart of the prediction process, along with alternative options to the benchmark method, *EstimateCellCounts2 (ECC2)*^42^. To that end, we utilized the CHILD cohort, with samples from cord blood at birth and peripheral blood at age 1 and 5 collected from approximately 800 children^11^. In addition, the absolute lymphocytes, monocytes, and granulocyte composition of the samples were quantified with a clinical-grade complete blood count (CBC), which was considered as the ground truth cell count measure for CHILD. We trained the optimal prediction pipeline in the age 5 samples and tested its validity in age 1 and cord samples. We further examined the performance of the optimized cell type prediction pipeline in independent validation cohorts, and compared them to the benchmark method *ECC2* in terms of mean absolute error (MAE) of predicted to CBC-based cell type proportions, whether the predicted proportions align with age-specific clinical intervals, and the amount of variance (adjusted R^2^) explained by the predicted proportions (**Figure 2**).

**Figure 2.**
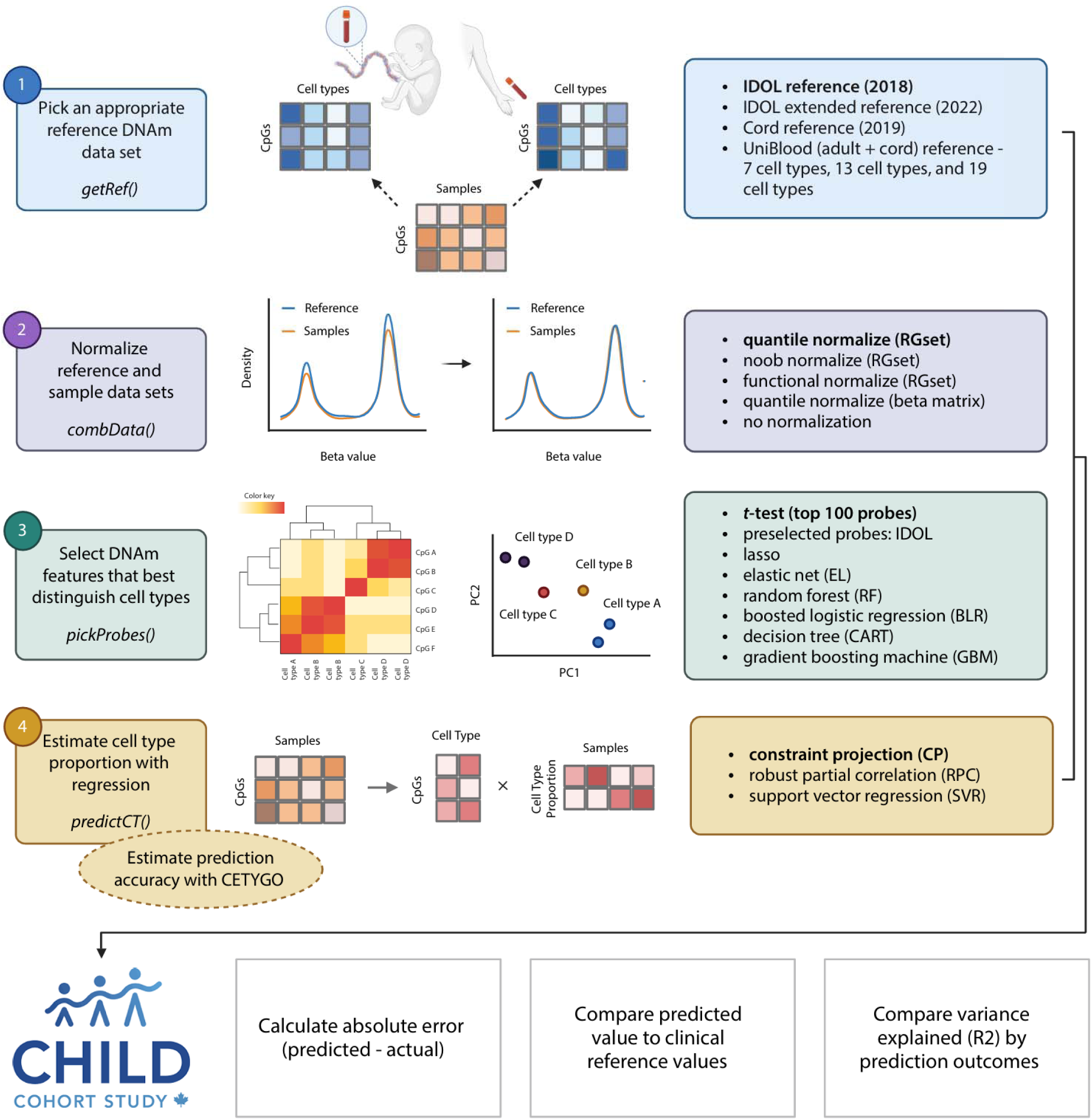
Schematic diagram of the standard reference-based DNAme cell type prediction pipeline and options evaluated in the optimization procedure. Top: Prediction steps and corresponding functions for implementation in bold (left) and options for the corresponding steps (right), with the bold option being the default in the benchmark method, *estimateCellCounts2,* and we used these default options as the starting point of the optimization process. Bottom: Methods for evaluation, including the calculation of absolute error, comparison to clinical ranges, and estimation of variance explained. Created with Biorender.com.

### Normalization method impacted the prediction performance across reference datasets

As the first step of the deconvolution process, we examined which blood reference dataset was most relevant to pediatric cohorts using CHILD age 5 samples. As no pediatric blood reference is available, we created three UniBlood references composed of both adult and cord blood cells to capture cellular heterogeneity across the developmental spectrum (**Table 1**). The UniBlood7 and UniBlood13 references considered the same cell types from cord and adult sorted cells to be the same population, aiming to identify development-agnostic cell-type DNAme signatures. In contrast, UniBlood19 considered cord and adult sorted cells as different populations (e.g. cord and adult monocytes), while taking into account the development-associated DNAme changes in immune cells.

**Table 1.** Cellular composition and dataset included for the three reference datasets we compiled in this study, UniBlood7, UniBlood13, and UniBlood19.

|  | UniBlood7 | UniBlood13 | UniBlood19 |
| --- | --- | --- | --- |
| FlowSorted.Blood.450k | Yes |  |  |
| FlowSorted.Blood.EPIC (IDOL) | Yes | only monocytes, neutrophils, and NK cells | only monocytes, neutrophils, and NK cells |
| FlowSorted.Blood.Extended.EPIC (Extended) |  | Yes | Yes |
| FlowSorted.Cord Blood.450k (Cord) | Yes | 6 randomly selected nRBC samples | 6 randomly selected samples for every cell type |
| Adult and cord cell types grouped? | Yes | No | No |
| Cell types included | **Seven cell types:** B cells, CD4+ T cells, CD8+ T cells, NK cells, monocytes, granulocytes, and nRBCs | **Thirteen cell types:** naïve and memory B cells, naïve and memory CD4+ T cells, naïve and memory CD8+ T cells, Tregs, NK cells, monocytes, neutrophils, basophils, eosinophils, and nRBCs | **Nineteen cell types:** neonatal, naïve, and memory B cells, neonatal, naïve, and memory CD4+ T cells, neonatal, naïve, and memory CD8+ T cells, adult Tregs, neonatal and adult NK cells, neonatal and adult monocytes, neonatal granulocytes, adult neutrophils, basophils, and eosinophils, and nRBCs |

Since the literature suggests the performance of the deconvolution to be dependent on normalization methods^37^, references and normalization methods were evaluated together. We compared six reference datasets (Cord^37^, UniBlood7, UniBlood13, UniBlood19, IDOL^42^, and Extended^39^; **Figure 1**, **Table 1**) in combination with normalization methods commonly employed for DNAme microarray data, including Quantile Normalization (Quantile), normal-exponential out-of-band normalization (Noob), and Functional Normalization (Funnorm), all applicable to *RGChannelSet* objects only^51^. To allow for applications in other data objects, we also assessed the performance of quantile normalization on beta matrices (Quantile.B) as well as the no normalization (None) conditions.

We compared the prediction performance of references by calculating the absolute error (AE) of predicted to lymphocyte, monocyte, and granulocyte proportions measured with CBC. We observed the performance of references to be dependent on the normalization method. For example, when comparing across references under Noob or Funnorm normalization, UniBlood13, UniBlood19, and Extended references resulted in the lowest AE (*p_adj_* < 0.05, **Figure 3**). On the other hand, IDOL reference yielded the lowest AE with Quantile normalization. For each condition, we also calculated the CETYGO score, which estimates deconvolution accuracy based on the deviation between a samples’ DNAme and the expected DNAme based the predicted cell type proportion and the reference profiles^48^. In concordance to the AE, Extended and UniBlood19 led to lower CETYGO score (indicating better prediction performance) with Noob and Funnorm normalization, whereas Extended and IDOL resulted in lower CETYGO score with Quantile normalization (**Supplementary Figure 1**). Additionally, Quantile normalization led to the lowest prediction error across the board for all reference panels (**Figure 3**). Statistically, with cell type and reference panel accounted for, the Quantile normalization method significantly outperformed all other methods (**Supplementary Table 1**). The RGChannelSet-based normalization methods consistently outperformed Quantile.B (*p_adj_* < 2.22e^-16^), which in turn significantly outperformed the no normalization condition (*p_adj_* = 1.68e^-5^). CETYGO score also corroborated the improved performance with Quantile normalization (**Supplementary Figure 1**). Despite the utility of CETYGO score, we showed that the metric failed to predict AE at the sample level (**Supplementary Figure 2**). Given the consistent performance of Quantile normalization, we decided to move forward with this normalization procedure.

**Figure 3.**
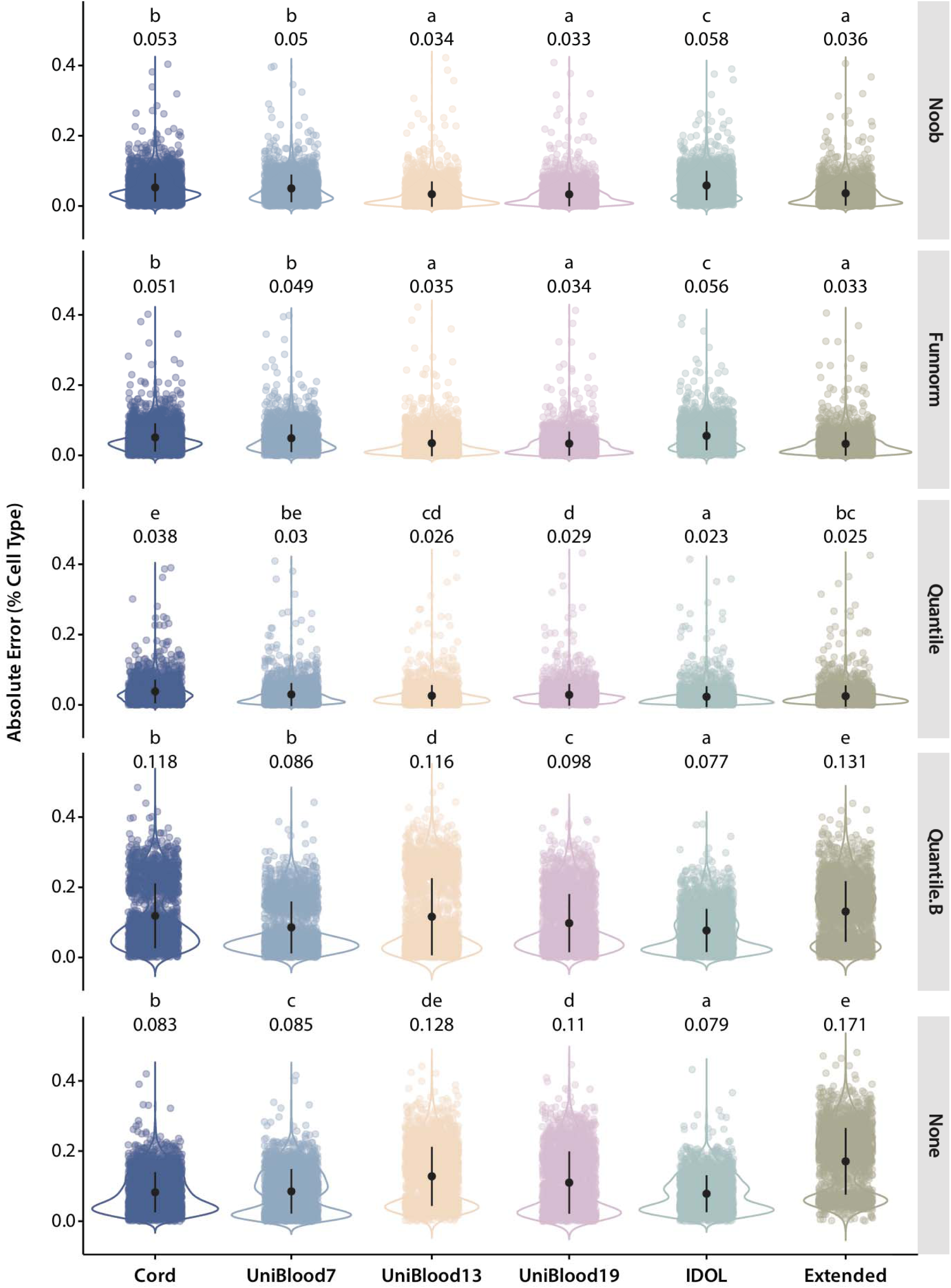
Violin plot of absolute prediction error of cell type proportions (predicted cell type proportion – actual proportion derived from complete blood count) across normalization methods (rows) and reference panels (columns) in CHILD age 5 samples. The mean absolute error of a given reference-normalization method pair condition is shown above the violin plot, with compact letter display indicating the significance of absolute error differences across reference panels within a normalization method. Significance is calculated with one-way ANOVA with post-hoc Tukey’s test.

### UniBlood19 reference unveiled epigenetic signatures associated with developmental immune cell transition

As the previous analysis focused on the prediction accuracy of the CBC subsets: lymphocytes, monocytes, and granulocytes, we next examined whether the predicted cell type proportions were accurate in additional cell type subsets (e.g. naïve CD4+ T cells). We compared the estimated proportions to age-specific clinical intervals reported in literature as we expected CHILD age 5 participants to align with healthy parameters^52–57^.

Using Quantile normalization, we observed a full alignment with clinical intervals for references predicting 6 to 7 cell types (Cord, UniBlood7, and IDOL) (**Figure 4A**). With increasing cell type resolution, we observed some deviation from the clinical intervals. Extended, UniBlood13, and UniBlood19 all overpredicted the proportion of CD8+ T cells and Tregs, and underpredicted the proportion of eosinophils (**Figure 4A**). Extended and UniBlood13 also overpredicted naive B cells, whereas UniBlood13 and UniBlood19 underpredicted memory CD4+ T cells. UniBlood13 reference yielded predictions that fell far outside the clinical intervals for multiple cell types, while the predictions with UniBlood19 and Extended were closer to expected based on clinical range. Noob normalization led to higher alignment with the clinical intervals for references predicting 12 or more cell types, whereas not performing normalization led to significant deviation from clinical intervals across references for most cell populations, highlighting the role of data normalization in deconvolution accuracy (**Supplementary Figure 3A-B**). Overall, IDOL reference had the highest percentage of predicted categories matching the clinical intervals across normalization methods, partly because it predicted fewer cell types (**Supplementary Table 2**), and Quantile normalization led to the highest alignment with clinical intervals, closely followed by Noob normalization.

**Figure 4.**
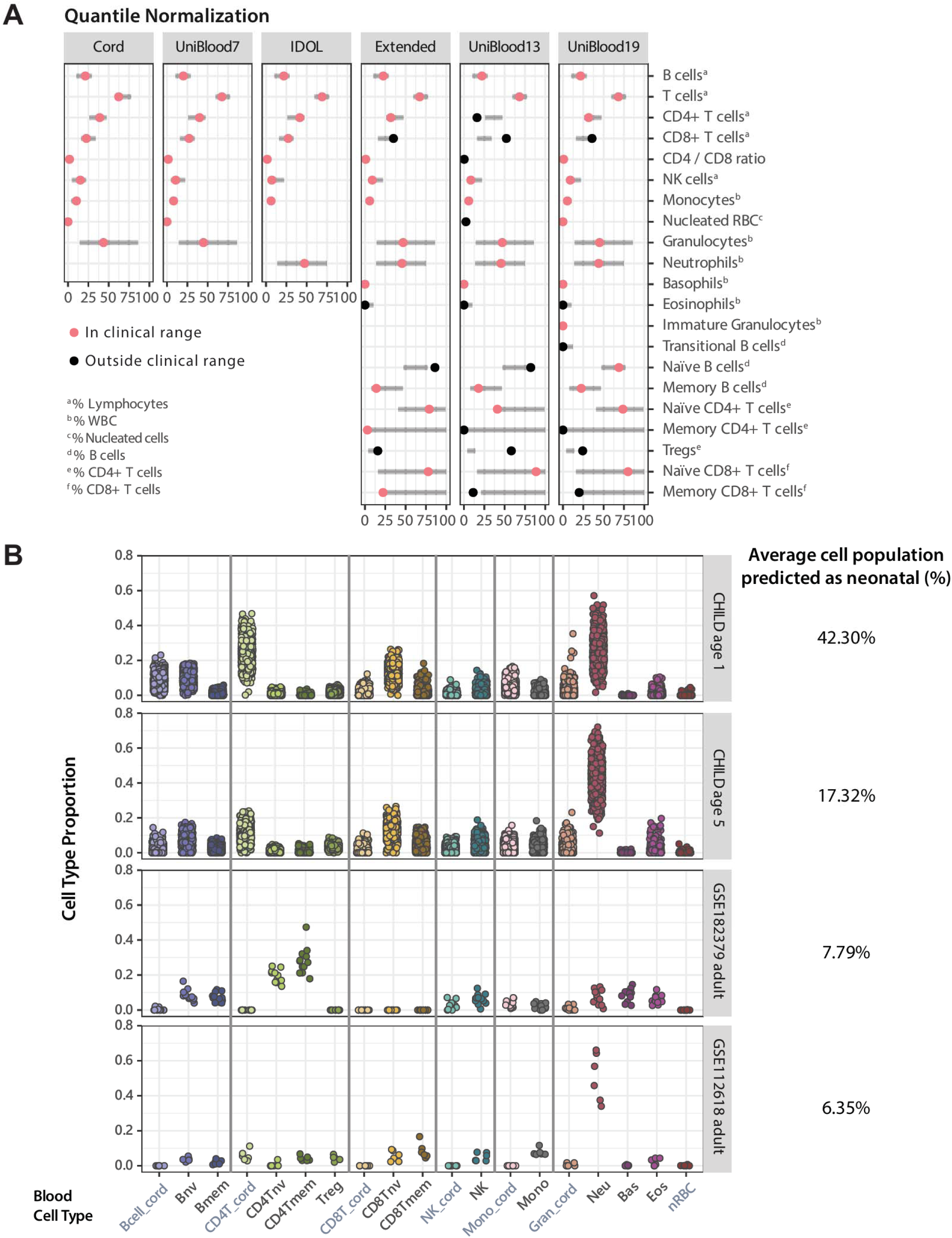
**A)** Age-specific clinical interval of hematological fractions reported in children at 5 years of age (gray bars) and predicted cell type proportions of CHILD age 5 blood samples based on DNAme profiles (dots). Six cell reference datasets were employed (columns) and Quantile normalization was performed. The clinical intervals were compiled from six publications with participants across sexes and of various genetic ancestry^1–6^, all measured with flow cytometry or hematology analyzers. The range presented here represents the most lenient 10^th^ and 90^th^ percentile range reported across literature (**Supplementary Table 3**). The colored dots are the median predicted proportion of a given cell population (salmon: inside clinical interval; black: outside clinical interval). For transitional B cells and immature granulocytes, the values correspond to estimated neonatal B cells and neonatal granulocytes proportions. WBC: white blood cells. **B)** Estimated blood cell proportions with the UniBlood19 reference panel, for four datasets with samples from difference age group: CHILD age 1, CHILD age 5, artificial mixture of adult-derived blood cells (GSE182379) and adult whole blood samples (GSE112618). The “_cord” suffix indicates predicted neonatal cells. nv: Naïve cells; mem: Memory cells; Treg: Regulatory T cells; NK: Natural killer cells; Mono: Monocytes; Gran: Granulocytes; Neu: Neutrophils; Bas: Basophils; Eos: Eosinophils; nRBC: Nucleated red blood cells.

By stratifying cord and adult immune cells in our reference panel, UniBlood19, we were able to delineate cell types in our pediatric samples that more closely resembled cord immune cells. These cell populations are more prevalent in peripheral blood during the pediatric time point, such as immature granulocytes (predicted as neonatal granulocytes) and transitional B cells (predicted as neonatal B cells)^16,54^. We predicted a low prevalence of both populations in CHILD age 5 samples, showing an underprediction of transitional B cells compared to the clinical interval (**Figure 4A**). To further illustrate the differences in deconvolution output across several reference datasets, we visualized the estimated cell type proportions of CHILD age 5 samples using the UniBlood19 reference (**Supplementary Figure 4**). We estimated a non-negligible proportion of neonatal cells, suggesting some of the immune cells at age 5 exhibit DNAme profiles more similar to that of neonatal cells. To explore whether this finding reflects biological changes during immune development or simply an artifact of the algorithm, we performed the same prediction with UniBlood19 reference for 3 other data sets: CHILD age 1, an artificial adult blood cell mixture (GSE182379) and an adult whole blood cohort (GSE112618). We observed higher estimates of neonatal cell type proportions in the pediatric age range and a decline with age (**Figure 4B**).

Finally, we wished to know which reference yielded predictions that explained the highest amount of variance. We observed that the deconvolution derived from UniBlood19 (the reference with the highest granularity) yielded the highest adjusted R2 estimates (**Supplementary Figure 5**). Overall, we selected two reference panels in our analyses: IDOL and UniBlood19. The IDOL reference was chosen for the lowest MAE when Quantile normalization was employed, and the UniBlood19 reference was selected due to the higher variance explained, and the potential of investigating cord to adult immune transition in future studies.

### Elastic net and random forest consistently performed well amongst feature selection methods

The next step in the deconvolution process is feature selection to identify a subset of DNAme loci best predicting each cell type. For the benchmark method *ECC2*, the default method for feature selection is based on top 100 probes selected from T-tests for each cell type. While several preselected probe sets for cell type deconvolution are available^39,42,58^, they fail to account for the normalization procedure, which can affect both the features selected and the coefficients estimated. We employed machine learning methods, including lasso, elastic nets (EN), random forests (RF), boosted logistic regression (BLR), category and regression tree (CART), and gradient boosting machine (GBM) to curate a list of DNAme probes predictive of cell type identity post normalization.

Similar to previous sections, we evaluated the prediction performance with MAE to CBC. Comparing the machine learning feature selection methods against the default method, t-tests using 1000 probes per cell type for prediction, we found multiple methods (all except for lasso and BLR) yielding comparable performance to t-tests for the IDOL reference (**Figure 5A**). In contrast, EN and RF significantly outperformed t-test and other methods for UniBlood19 reference (**Figure 5B**). To assess the effect of selected CpGs on cell type proportion prediction performance, we altered the number of probes supplemented for each cell type (*k)* from 10 to 5000. No clear relation between *k* and MAE was observed for IDOL reference, but t-test, EN, and RF consistently resulted in low MAE, with the first two yielding slight increased MAE past *k* = 1000, whereas RF performance improved with increasing *k* (**Figure 5C**). For the UniBlood19 reference, MAE decreased as *k* increased for t-test, RF, and EN, with slight exception with *k* = 5000 (**Figure 5D**). Specifically, EN consistently yielded the lowest MAE with *k* > 300. Overall, EN with *k* = 1000 performed well for both IDOL and UniBlood19 references, and was chosen as the feature selection method moving forward. Finally, we observed an equivalent performance across regression methods in the CHILD blood sample, so we decided to move forward with constraint projection that is well adopted in blood (**Supplementary Figure 6**)^36^.

**Figure 5.**
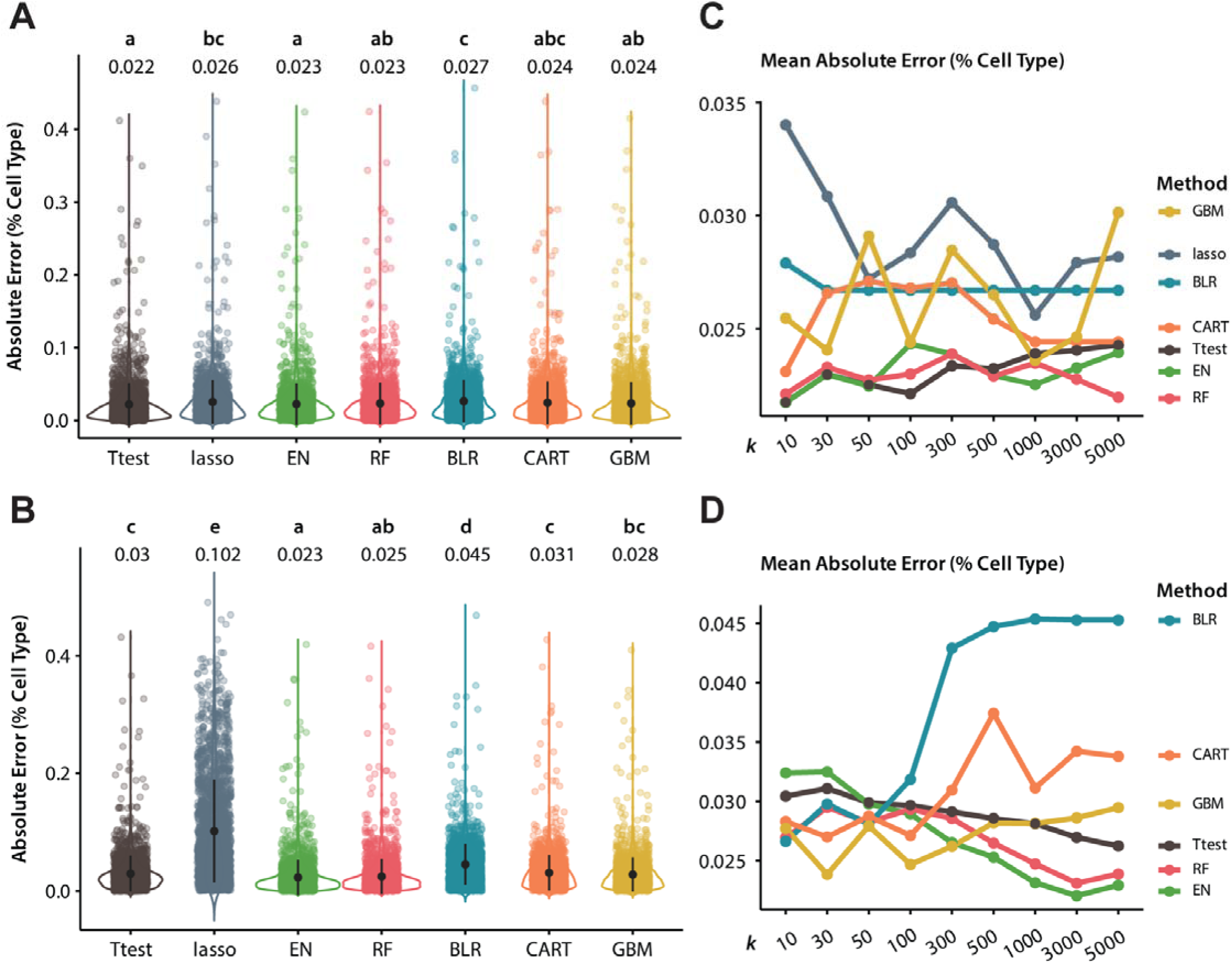
**(A-B)** Absolute prediction error of cell type proportions (predicted – true) across machine-learning-based feature selection methods based on **A)** IDOL reference and **B)** UniBlood19 reference with Quantile normalization, using top 1000 cell-type-specific probes per cell type. **(C-D)** MAE of predicted cell type proportions across varying number of available features (*k*) for each feature selection method based on **C)** IDOL reference and **D)** UniBlood19 reference

### The optimized pipeline outperformed benchmark methods in test datasets

After optimizing the pipeline in CHILD age 5 samples, with quantile normalization, IDOL or UniBlood19 reference, EN feature selection, and constraint projection (hereafter referred to as the *CellsPickMe* algorithm), we evaluated the *CellsPickMe* performance by comparing it against benchmark algorithms in the reserved CHILD and external validation cohorts. We first explored the performance of CHILD age 1 peripheral blood and cord blood. We found the *CellsPickMe* algorithm with UniBlood19 reference led to lower MAE than *ECC2* with either the IDOL or Extended reference (**Figure 6A**). We observed a similar phenomenon in the CHILD cord blood samples as well. Although not an exclusive cord blood reference, CellsPickMe with the UniBlood19 reference achieved lower MAE than *ECC2* with the Cord reference, with or without using the preselected IDOL probes (**Figure 6B**). Furthermore, the *CellsPickMe* algorithm predicted predominantly neonatal cells with comparable estimated nRBC proportions to using the Cord reference, demonstrating the algorithm’s ability to detect DNAme signature specific to neonatal cells and applied them in deconvolution (**Figure 6C**).

**Figure 6.**
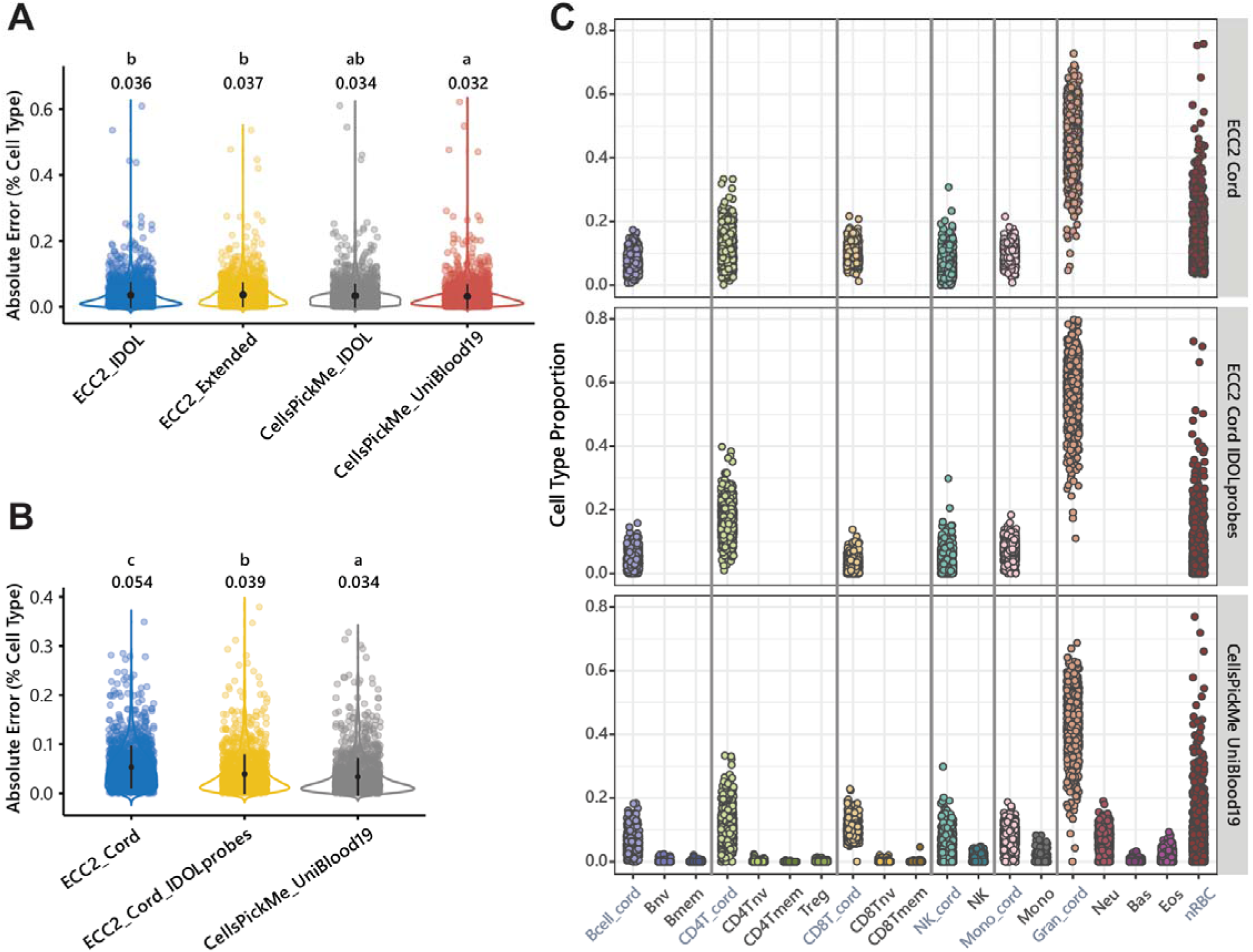
**(A-B)** Absolute prediction error of cell type proportions in the *CellsPickMe* optimized deconvolution algorithm and the benchmark estimateCellCounts2 method in reserved CHILD **A)** age 1 and **B)** cord blood samples. **C)** Deconvolution prediction results for CHILD cord blood using either the benchmark estimateCellCounts2 method, with or without the IDOL probe list, or CellsPickMe with UniBlood19 reference.

Finally, we validated the performance of *CellsPickMe* in an independent cohort. We utilized both the IDOL and UniBlood19 references and compared their performance to *ECC2* using either the IDOL or Extended reference. In GSE112618 (adult whole blood), we observed the lowest MAE using CellsPickMe with IDOL (*p_adj_*< 0.05), and a comparable MAE between CellsPickMe with UniBlood19 reference and *ECC2* with Extended reference (**Figure 7**). Additionally, CellsPickMe with IDOL significantly outperformed all other methods in adult blood, demonstrating the utility of machine learning-based feature selection method even with the same reference dataset being employed.

**Figure 7.**
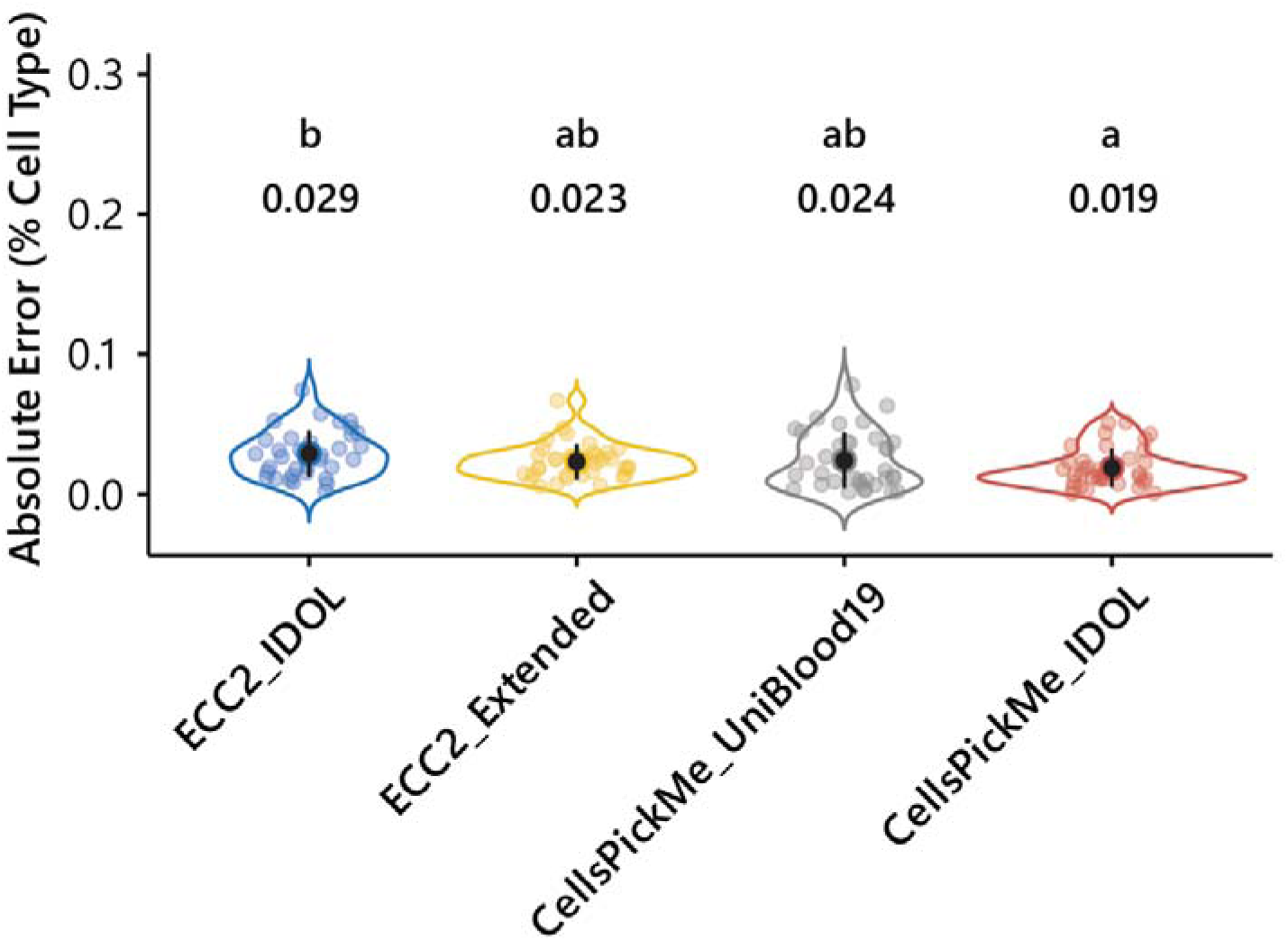
Absolute prediction error of cell type proportions in the CellsPickMe optimized deconvolution algorithm and the benchmark estimateCellCounts2 method in an external validation adult whole blood dataset (GSE112618).

## Discussion

Through systematic and rigorous evaluation of algorithmic performance, we optimized a cell deconvolution procedure, *CellsPickMe*, that can be applied to blood datasets across the developmental spectrum. The algorithm enhanced cell type prediction with features such as multi-age reference datasets, data-driven feature selection, and performance evaluation using CETYGO. In our optimization process, we noted that quantile normalization with the IDOL reference led to low prediction error, whereas the UniBlood19 reference uniquely elucidated cell type proportion differences in longitudinal birth cohort studies with matched cord and pediatric samples. Machine learning approaches, like EN and RF, not only matched or slightly improved prediction accuracy, but also offered a defined set of cell type-specific probes for downstream analysis. Ultimately, the optimized pipeline — featuring quantile normalization, the UniBlood19 or IDOL reference, EN, and constraint projection — surpassed the existing benchmark in predicting cell types for CHILD age 1 and cord blood samples, as well as external adult datasets.

We were inspired to create the UniBlood reference datasets because of one curious phenomenon observed in the PCA of sorted cells’ DNAme, that there was a distinct maturity axis amongst the top PCs. Interestingly, the cord samples exist on this continuum as an extension of the naïve cell state. When we overlayed DNAme profiles of various developmental time points with that from the sorted cells, there was a distinct cluster of age 1 samples away from the cord blood, and another of the age 5 samples that was closer to the adult blood than their age 1 counterpart. This suggests pediatric cell types have intermediate DNAme profiles not currently accounted for in existing references. Our findings align with immune phenotype profiling with mass cytometry, which has shown that the immune landscape changes drastically immediately post-birth, with uncorrelated profiles between cord blood to that one-week post birth.^15^ We also observed this rapid shift in the immune system reflected in the optimization process, where the IDOL reference consistently showed high prediction accuracy and high alignment with age-specific clinical intervals. This suggests that, while the DNAme profiles of pediatric blood at age 5 is distinct from both cord blood and adult blood, it is more like the latter. The observation speaks to the need to have a reference dataset with unified cord and adult blood cells to accurately assess this cellular transition in the pediatric age range.

With the postnatal loss of nRBCs and rapid shift in neonatal immune system, DNAme research has reported challenges in integrating cord blood and pediatric peripheral blood^33,34^. Applying adult references on cord blood has been shown to be inadequate in accounting for cellular heterogeneity^37^. Similarly, we have demonstrated that applying cord references in pediatric peripheral blood samples yielded poor performance. The three UniBlood references offer the strategical advantage of harmonizing deconvolution of samples collected across multiple developmental time points under a single framework. This facilitates comparisons of association test results across developmental time points because cellular composition can be accounted for with the same constituent cell types. We also noted that UniBlood7 and UniBlood13 underperformed compared to UniBlood19, potentially because the former two references intended to identify developmentally agnostic cell type markers when cell-type-specific DNAme patterns changed significantly with early development. As a result, UniBlood19, which treated cord and adult blood cells as independent identities, was able to better capture cell-type-specific DNAme signatures and resulted in improved prediction performance.

The literature has shown that neonatal immune cells are distinct from their adult counterpart in both identity and functions, including B cells^59–61^, T cells^62–64^, monocytes^65,66^, NK cells^67,68^, and granulocytes^69,70^. The neonatal immune system needs to be simultaneously capable of tolerating the maternal environment, as well as rapidly adapting to the pathogen exposure upon birth.^59,63,64^ Because of the rapid shift from maternally derived immunity after birth, it has been observed that immature granulocytes and transitional B cells are more prevalent in cord blood, with diminishing proportions later in life.^60,53,54,15,69,16^ The neonatal immune cells’ DNAme profile was sufficiently unique that *CellsPickMe* was able to accurately reproduced this dynamic, predicting high proportion of neonatal immune cells in cord blood and decreasing proportion in samples of older ages. Furthermore, the UniBlood19 reference can estimate neonatal cell populations, such as neonatal B cells and neonatal granulocytes. Quantifying these cell types not only informs immune development but also functions as relevant medical biomarkers. For instance, as transitional B cells serve as a pivotal link between immature and mature B cells – negative selection against autoreactive clones – altered frequency of transitional B-cell subsets have been linked to systemic lupus erythematosus and other autoimmune disorders^71–73^. In the example of immature granulocytes, we hypothesize that the captured DNAme signature is linked to not just immature granulocytes, but also low density neutrophils (LDNs), as their identity overlap in cord blood^69^. LDNs are lighter neutrophils that are found in peripheral blood mononuclear cells after gradient centrifugation^74^. These cells possess immunosuppressive or proinflammatory characteristics, and have higher abundance in cord blood, pregnancy, or in patients with autoimmune diseases, cancer, and infection^21,74–77^. Our findings suggest that epigenetics is a viable avenue to explore immune cell development and transition across developmental trajectory, supplementing useful yet distinct information to commonly studied immune markers like cell surface markers and cytokine milieu.

As cell identity and epigenetics are inextricably linked, one of the challenges was the selection of DNAme sites that are most predictive of given cell types. We aimed to build an algorithm that is both robust and adaptive. Instead of generating a curated list of DNAme sites, we performed the feature selection step post-normalization because the prediction performance is dependent on the preprocessing pipeline, as evidenced by both the discrepancy in MAE and alignment with clinical interval with and without normalization. We explored a handful of machine learning methods with embedded regularization steps to identify the probe sets that best predict cell type identity.

Surprisingly, the benchmark method of selecting 100 probes with t-tests consistently performed well across experimental conditions. The results point to the relevance of cross-validating a range of machine learning algorithms for optimization purposes. The observation echoed the “no free lunch” theorem for optimization, which proposes that the optimal solution of a given problem varies, and no one solution is superior for all problems or data sets.^78,79^

We also tested a range of *k* — number of initial features available for the regularization algorithm — and showed, for IDOL reference, optimal performance can be achieved with t-tests, EN, and RF, with as few as 10 probes per cell type. This option can potentially be implemented for users who wish to improve deconvolution efficiency both in terms of time and computational memory usage. Ultimately, we observed that EN and RF consistently resulted in low prediction error at higher *k*. For users who intend to investigate the biological process underlying immune cell-specific DNAme signature, we recommend performing feature selection with EN or RF with *k* between 1000 and 5000.

While our study hints towards the application of DNAme-based deconvolution in studying immune system dynamics, there are some aspects that warrant further consideration. For one, CBC was considered as the ground truth cell count measure for CHILD. Despite being a well-established method for examining cellular composition in clinical settings, it cannot resolve lymphocyte subsets, which is critical for understanding immune responses^80^. Furthermore, collapsing the subsets can potentially bias the performance assessment process. We also do not have the genetic information of the samples included in the reference datasets. There can potentially be cell-type-specific methylation quantitative trait loci that drive feature selection, which can impact the deconvolution performance. Another consideration when utilizing *CellsPickMe* is the relatively longer runtime compared to other methods, but this is due to the inclusion of normalization and pre-selection of the prediction features which increases the rigor and reproducibility of our tool. While our method is more time-consuming, we have shown that these two steps significantly improve deconvolution outcomes, but computational demands should be considered when approaching these methods. Finally, despite having validated the prediction accuracy of *CellsPickMe* in CHILD as well as external cord and adult blood datasets, we have not examined its performance in more diverse settings, such as non-predominantly European genetic ancestry, elderly populations, or diseased conditions. Future studies can address the gap by comparing CellsPickMe predicted cell type proportions to empirical cell counts in these cohorts.

In this study, we demonstrated *CellsPickMe*’s superior performance beyond pediatric datasets, to cord and adult blood samples as well. Additionally, the feature selection methods employed in *CellsPickMe* demonstrated improved prediction performance across validation datasets we tested and output a list of cell-type-specific DNAme sites that can be further explored. We also reported on the utility of applying CETYGO score for evaluation of the appropriateness of given reference datasets and normalization methods and incorporated this internal error assessment metric in the package for easy application. Overall, *CellsPickMe* improved upon existing cellular deconvolution methods in blood, enhancing researchers’ ability to explore and account for immune dynamics in their DNAme investigations, especially in longitudinal pediatric studies.

## Methods

### Cohort description

a. Canadian Healthy Infant Longitudinal Development (CHILD)

The CHILD cohort is a population-based prospective birth cohort study that recruited 3621 pregnant mothers across four sites in Canada^11^. At birth, the cord blood samples of children enrolled in the study were collected from the umbilical cords (n = 838). At age one and five years (referred to as age 1 and age 5 hereafter), peripheral blood samples were also collected as previously described (n = 1616)^81^.

Out of the participants, the majority of them identifies as White (75%), followed by East Asian (8%), South East Asian (5%), First Nations (3%), Multiracial (3%), South Asian (2%), Black (2%), and Hispanic (2%). DNA was extracted from the cord, age 1, and age 5 blood samples, and the DNAme profiles assayed.

b. Isle of Wight (IOW)

The IOW birth cohort is a whole population prospective study that enrolled all children born on the Isle of Wight, UK, between January 1989 and February 1990 (n = 1536)^13^. Blood samples were collected from the participants at birth (n = 777) as Gutherie cards, and at age 10 (n = 406), 18 (n = 141), and 26 years (n = 295) as peripheral blood draws. The DNAme profiles were then assessed as previously described^82^.

### DNA sample collection and methylation array for the CHILD cohort

DNA was isolated from cord and peripheral whole blood from CHILD cohort samples using the DNeasy Blood & Tissue Kit (Qiagen, Venlo, Netherlands). The DNA concentration and quality were evaluated via a NanoDrop 8000 Spectrophotometer (Thermo Fisher Scientific, Waltham, MA, USA). For DNA methylation profiling, the purified DNA was bisulfite converted with the EZ-96 DNA Methylation Kit (Zymo Research, Irvine, CA, USA) and assayed with the Infinium MethylationEPIC BeadChip array (Illumina, San Diego, CA). The resulting raw intensity IDAT files contained 866,836 data points covering 863,904 CpG sites.

### CHILD cohort DNA methylation data processing

All available CHILD whole blood DNAme data were read into and further processed with RStudio (version 4.0.3).^83^ Quality control was performed using the *ewastools* v1.7 package based on technical parameters evaluated on the 636 control probes^84^. The *minfi* v1.44 package was then used to evaluate methylated and unmethylated intensities, and check for sex concordance between reported and predicted biological sex^85^. Outliers and poorly performing samples were then identified, with the former detected with the *lumi* package^51,86^, and the latter defined as those with a high detection p-values (*p* > 0.01 in > 1% of probes), DNAme intensities significantly deviating from the average of negative control probes, or a bead count of less than three on more than 1% of probes. The 59 SNP probes on the array were used to further confirm sample relatedness amongst the cord blood, age 1 and age 5 samples for each child. Samples that failed one or more of the quality control metrics were removed (n = 29). After quality control, technical replicates were also removed (n = 22). Overall, 807 samples at age 5 and 795 samples at age 1 remained after sample filtering for cell deconvolution.

### Pan-age blood reference-based cell deconvolution

We followed the well adopted statistical procedures in cell type deconvolution^36^, as outlined in **Figure 2**. The method assumed the statistical model:

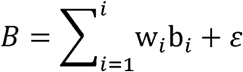

where B represented the beta matrix of sample DNAme, and w and b represented the proportion and beta value profile of cell type *i* in the sample, respectively. The error term represented variability from either cell types not included in the reference dataset, other biological sources of DNAme variability, or technical noise. The prediction algorithm aimed to calculate b_i_ in a subset of cell-type-specific DNAme sites in the reference dataset for the estimation of w_i_ in the sample dataset.

#### Step 1. Reference selection

To create a cell type deconvolution algorithm that allows for cell type prediction in both cord and peripheral blood, four published DNAme datasets of purified blood cell types were combined to create the UniBlood references – the *FlowSorted.Blood.450k* (Reinius) ^87^, *FlowSorted.Blood.EPIC* (IDOL)^38^, *FlowSorted.CordBlood.450k* (Cord)^37^, and *FlowSorted.Blood.Extended.EPIC* (Extended)^39^ references. The first two of these encompass 6 immune cell types (B-cells, CD4+ T-cells, CD8+ T-cells, natural killer [NK] cells, monocytes, and granulocytes), whereas the cord blood reference also includes nRBCs. Finally, the Extended reference^39^ includes 12 cell subtypes sorted at finer granularity (naïve and memory B cells, naïve and memory CD4+ T cells, naïve and memory CD8+ T cells, regulatory T cells (tregs), natural killer (NK) cells, monocytes, basophils, eosinophils, and neutrophils). We curated three UniBlood references, with different dataset composition.

1. UniBlood7: Included IDOL, Reinius, and Cord reference datasets. The cord and adult blood cells of the same types were grouped together (e.g. CD4+ T cells of the 3 datasets were grouped together and considered as one cell type).
2. UniBlood13: Included Extended and nRBCs from the Cord reference. As nRBCs have the most distinct DNAme profile and is absent in adult blood, this data set assumed the other cell types maintain similar DNAme profiles across development, and only the absent nRBC is required to deconvolute cord and pediatric samples.
3. UniBlood19: Included Extended and Cord reference datasets. This reference set considers neonatal and adult blood cells as distinct entities. This data set assumed that neonatal and adult blood cells’ DNAme profiles are distinct. To ensure equal representation of cord and adult cell types, we randomly sampled 10 samples for each cell type in the Cord reference (seed = 1234) prior to combining the datasets.

We compared the influence of UniBlood references on deconvolution accuracy, against that of the Cord, IDOL, and Extended reference.

#### Step 2. Normalization of reference and sample datasets

Normalization aimed to reduce batch effect between reference and sample datasets and generate comparable sample distribution to ensure the estimated coefficients in the reference applies to the samples. We compared common normalization strategies, including those applicable to RGChannelSet object (Noob normalization [Noob], as implemented with minfi::preprocessNoob, Functional normalization [Funnorm], as implemented with minfi::preprocessFunnorm, Quantile normalization [Quantile], as implemented with minfi::preprocessQuantile), those applicable to beta matrix (Quantile normalization [Quantile.B], as implemented with limma::normalizeQuantiles), and a no normalization condition.

#### Step 3. Feature selection

The process of reference-based cell type prediction then selects probes that best discern the cell types included in the reference. The benchmarking approach uses T-tests to select for probes with the highest mean difference when comparing a given cell type with all others. Alternatively, pre-selected probe sets exist, including IDOL probes^42^, IDOL-ext probes^39^, and DNAase hypersensitive sites (DHS)^88^. We proposed new probe selection methods based on machine learning algorithm that has intrinsic feature selection process. Random forest (RF), elastic net (EN), boosted logistic regression (BLR), gradient boosted machine (GBM), and classification and regression tree (CART) were implemented with leave-one-out cross validation (LOOCV) using the caret package.

#### Step 4. Coefficient estimation and regression-based prediction

Sample cell type proportions are estimated with linear regression under constraints such that w*_i_* ≥ 0 and ∑*^i^_i=1_* w*_i_* ≤ 1. The most common implementation of such conditions is constraint projection (CP) with quadratic programming to optimize the two inequality constraints (Houseman)^89^. Two other popular approaches are robust partial correlation (RPC) and support vector regression (SVR) (EpiDISH & CIBERSORT)^90^. These two methods apply the normalization constraints *a posteriori* by setting negative estimates to 0 and readjusting each estimate proportionally so that the cell types sum up to 1. We compared the deconvolution performance with the three regression methods.

### Model Performance Assessment

To compare the performance across conditions for a given prediction step, three main metrics were assessed: absolute error (AE), comparison with age-specific clinical count interval, and variance explained accounting for the number of covariates (adjusted R^2^) for DNAme. For the CHILD samples, the complete blood count (CBC), comprised of lymphocytes, monocytes, and granulocytes portions, was considered as the ground truth. After inference, the predicted proportions were summed for each of the three cell subsets, with neutrophils, basophils, and eosinophils adding up to the granulocyte proportion, and T cells, B cells, and NK cells adding up to the lymphocyte proportion. AE was calculated as the difference between predicted and actual cell type proportions for each subset. One-way ANCOVA tests were applied across conditions to evaluate the differences in AE distribution, with post-hoc Tukey’s test to correct for multiple testing and identify the groups with significantly different distributions. Additionally, the CEll TYpe deconvolution GOodness (CETYGO) score was calculated as the root mean squared error of the DNAme based on the estimated cell type proportions and the observed DNAme^48^. The CETYGO score ranges from 0 to 1, with a lower score indicating a better fit of the reference dataset and prediction procedure for the samples of interest.

For the comparison with the clinical interval, we calculated the median predicted proportion for each combination, and considered the prediction to overlap with the clinical range reported in the literature if the median falls within the 10^th^ and 90^th^ percentile of the interval.^16,52,53,55–57^ Finally, we assessed the variance in the DNAme data explained by cell type proportion estimated under across reference datasets and normalization methods. The estimated proportions were first summarized with principal component analysis. We applied EWAS on the estimated cell type PCs accounting for 90% of variance explained in the estimated proportion.

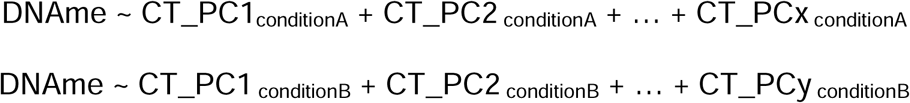

We then compared the distribution of the adjusted R^2^ across measured DNAme sites among prediction conditions. One-way ANOVA with post-hoc Tukey’s test were utilized to calculate statistical significance.

### Code Availability

An R package is available on GitHub (https://github.com/maggie-fu/CellsPickMe) and can be installed through devtools::install_github(“maggie-fu/CellsPickMe”).

## Acknowledgments

We would like to acknowledge the dedicated and tremendous work by Tanya Erb that enabled data access, facilitated project administration, and enabled inter-lab collaboration. We are also grateful to our colleague Beryl Zhuang, for establishing and coordinating smooth collaborations. We would also like to thank the CHILD Cohort Study (CHILD) participant families for their dedication and commitment to advancing health research. Furthermore, we would like to acknowledge the work of Meghan Azad as a CHILD site leader for implementing the study and making our work possible. CHILD was initially funded by CIHR and AllerGen NCE. Visit CHILD at childstudy.ca for more information.

## Author contributions

Conceptualization: MPF, MSK

Methodology: MPF, EIND, SMM

Formal Analysis: MPF, KE

Investigation: MPF, KE, EIND

Resources: NK, PM, ES, PS, TJM, JWH, SET, MSK

Visualization: MPF

Funding acquisition: MSK

Supervision: MSK

Writing – original draft: MPF

Writing – review & editing: MPF, KE, EIND, SMM, CK, NK, JWH, MSK

