## Supplementary Figure 1 for "Advancing Pediatric and Longitudinal DNA Methylation Studies with CellsPickMe, an Integrated Blood Cell Deconvolution Method"

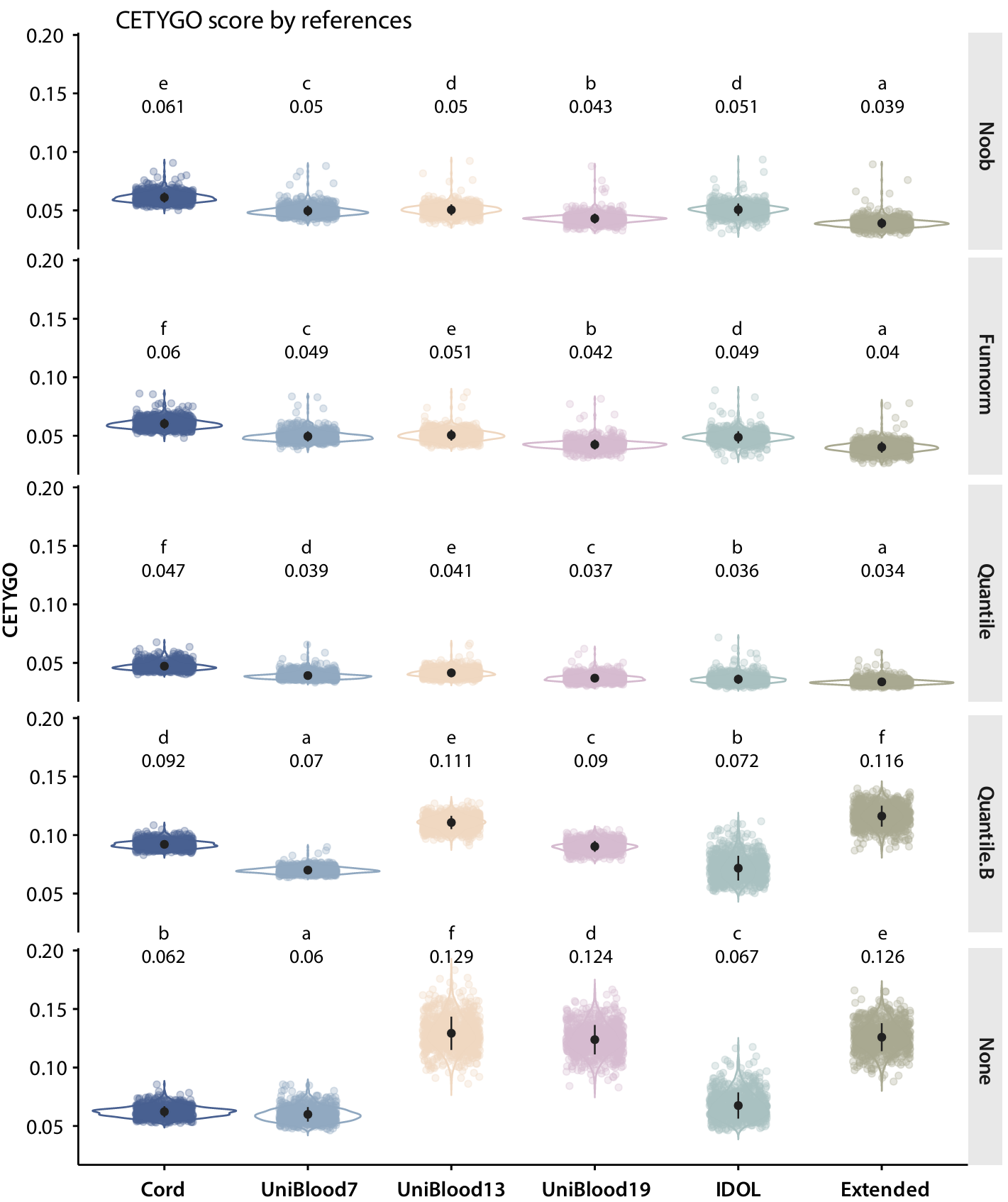

**Supplementary Figure 1.** Violin plot of CEYGO score across normalization methods (rows) and reference panels (columns) in CHILD age 5 samples. The mean CETYGO score of a given reference panel-normalization method pair condition is shown above the violin plot, with compact letter display indicating the significance of absolute error differences across reference panels within a normalization method. Significance is calculated with one-way ANOVA with post-hoc Tukey’s test, with multiple test correction with Bonferroni method.
