## Supplementary Figure 2 for "Advancing Pediatric and Longitudinal DNA Methylation Studies with CellsPickMe, an Integrated Blood Cell Deconvolution Method"

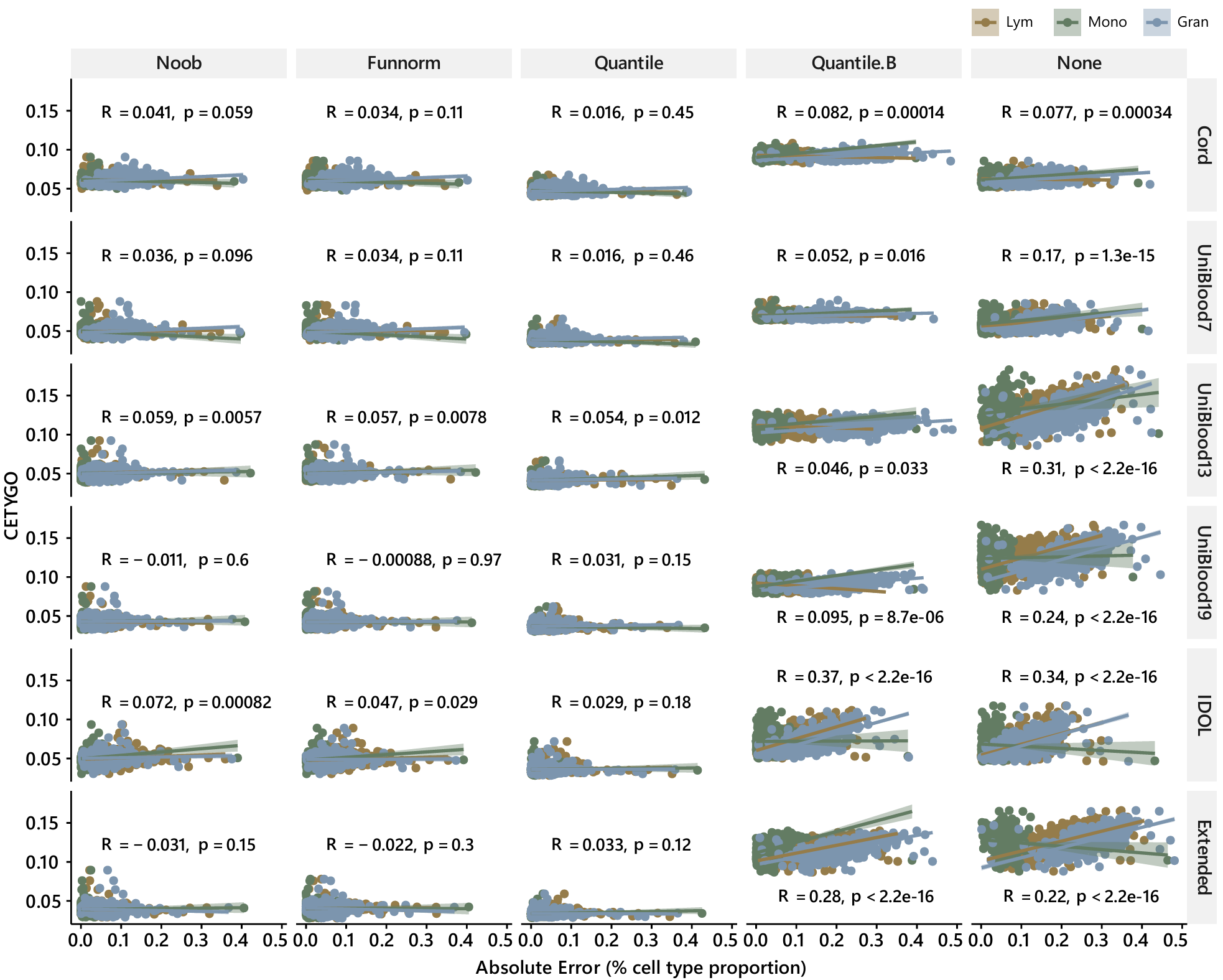

**Supplementary Figure 2.** Correlation of absolute prediction error of cell type proportions (predicted cell type proportion – complete blood count proportion) and CETYGO score for age 5 CHILD samples for lymphocytes (brown), monocytes (green), and granulocytes (blue).
