## Supplementary Figure 3 for "Advancing Pediatric and Longitudinal DNA Methylation Studies with CellsPickMe, an Integrated Blood Cell Deconvolution Method"

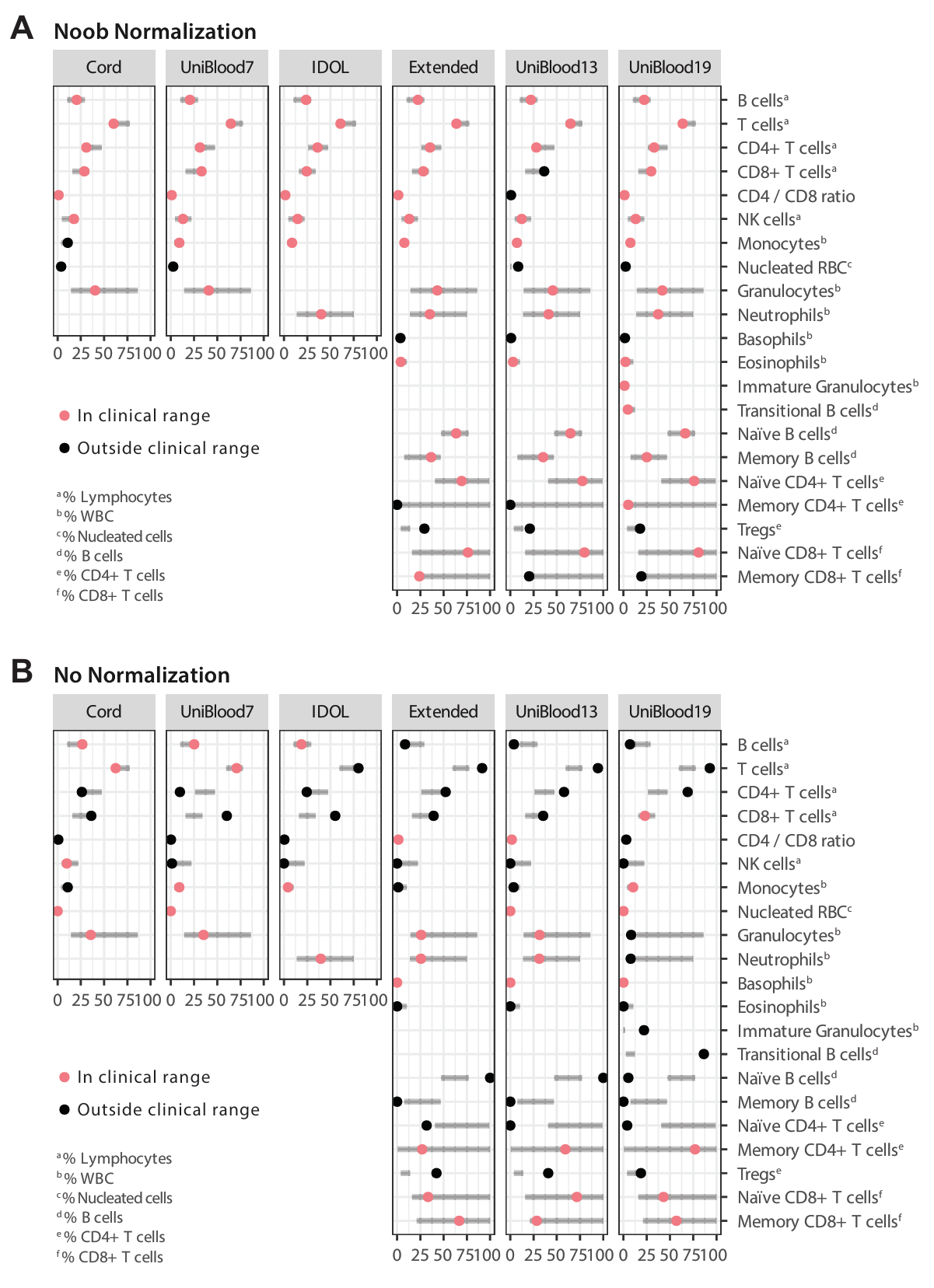

**Supplementary Figure 3.** Age-specific clinical interval of hematological fractions reported in children at 5 years of age (gray bars) and predicted cell type proportions of CHILD age 5 blood samples based on DNAme profiles (dots). Six cell reference datasets were employed (columns) where **A)** Noob or **B)** no normalization was performed. The clinical intervals were compiled from six publications with participants across sexes and of various genetic ancestry^1–6^, all measured with flow cytometry or hematology analyzers. The range presented here represents the most lenient 10^th^ and 90^th^ percentile range reported across literature (Supplementary Table 3). The colored dots are the median predicted proportion of a given cell population (salmon: inside clinical interval; black: outside clinical interval). For transitional B cells and immature granulocytes, the values correspond to estimated neonatal B cells and neonatal granulocytes proportions. WBC: white blood cells.
