## Supplementary Figure 4 for "Advancing Pediatric and Longitudinal DNA Methylation Studies with CellsPickMe, an Integrated Blood Cell Deconvolution Method"

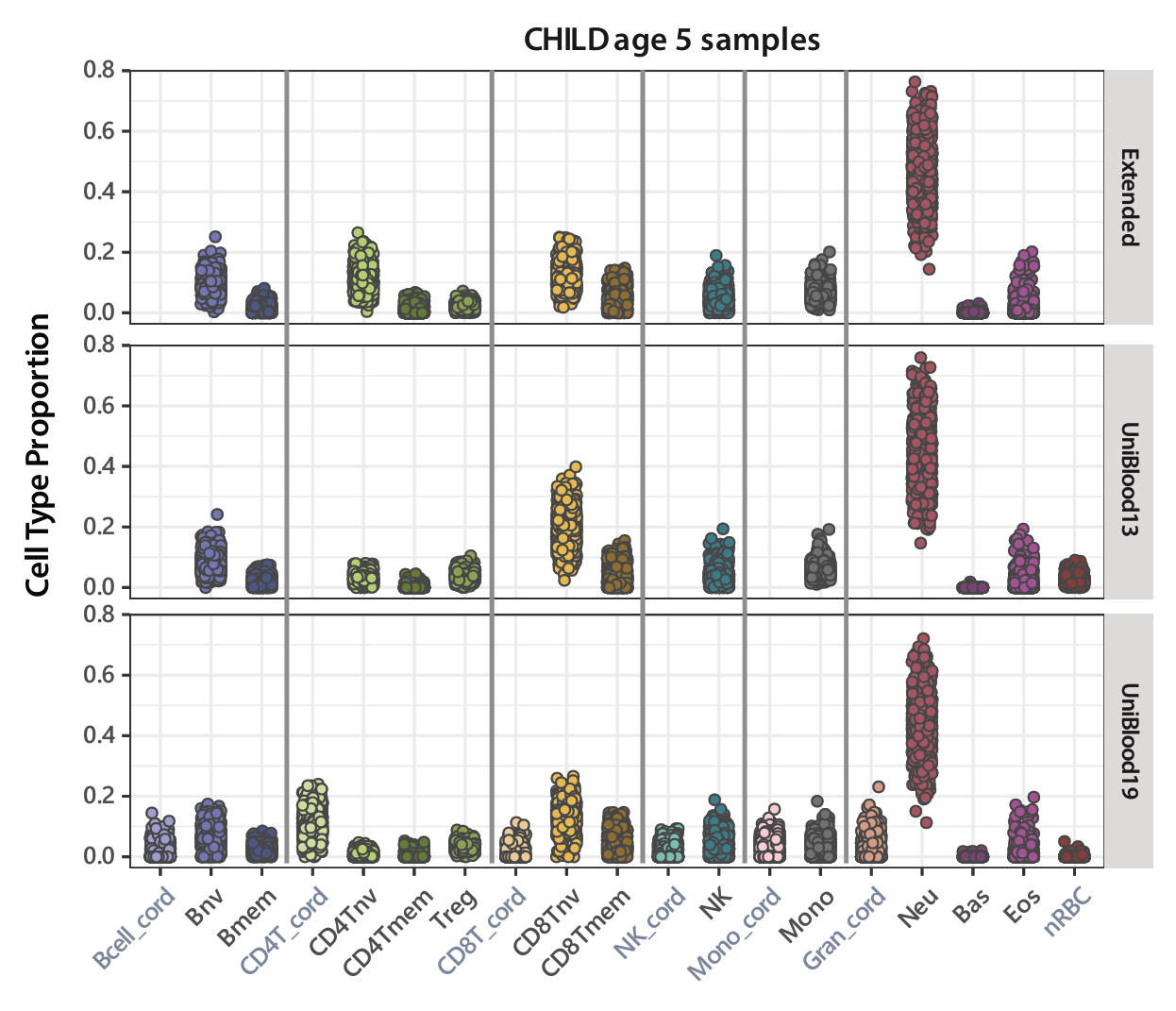

**Supplementary Figure 4.** Estimated blood cell proportions in age 5 CHILD samples across Extended, UniBlood13, and UniBlood19 reference panels. The “_cord” suffix indicates predicted neonatal cells. nv: Naïve cells; mem: Memory cells; Treg: Regulatory T cells; NK: Natural killer cells; Mono: Monocytes; Gran: Granulocytes; Neu: Neutrophils; Bas: Basophils; Eos: Eosinophils; nRBC: Nucleated red blood cells.
