## Supplementary Figure 5 for "Advancing Pediatric and Longitudinal DNA Methylation Studies with CellsPickMe, an Integrated Blood Cell Deconvolution Method"

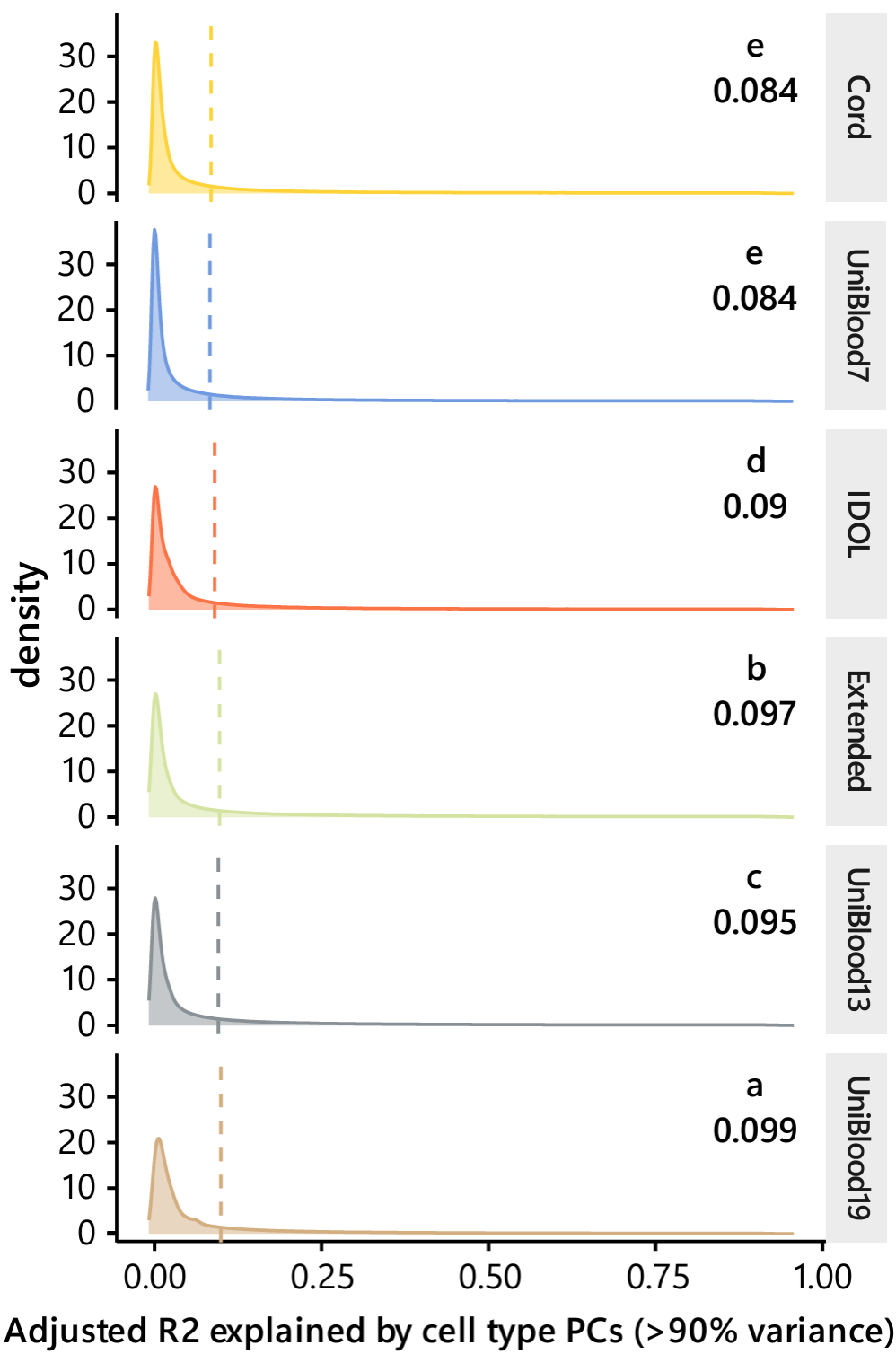

**Supplementary Figure 5.** Density plot showing the distribution of adjusted R^2^ as DNAme for each DNAme is fitted in a regression model with cell type estimates with each reference panel. The vertical dotted line corresponds to the mean of the density distribution. Significance is calculated with one-way ANOVA with post-hoc Tukey’s test, with multiple test correction with Bonferroni method, and shown with compact letter displayed.
