## Supplementary Figure 6 for "Advancing Pediatric and Longitudinal DNA Methylation Studies with CellsPickMe, an Integrated Blood Cell Deconvolution Method"

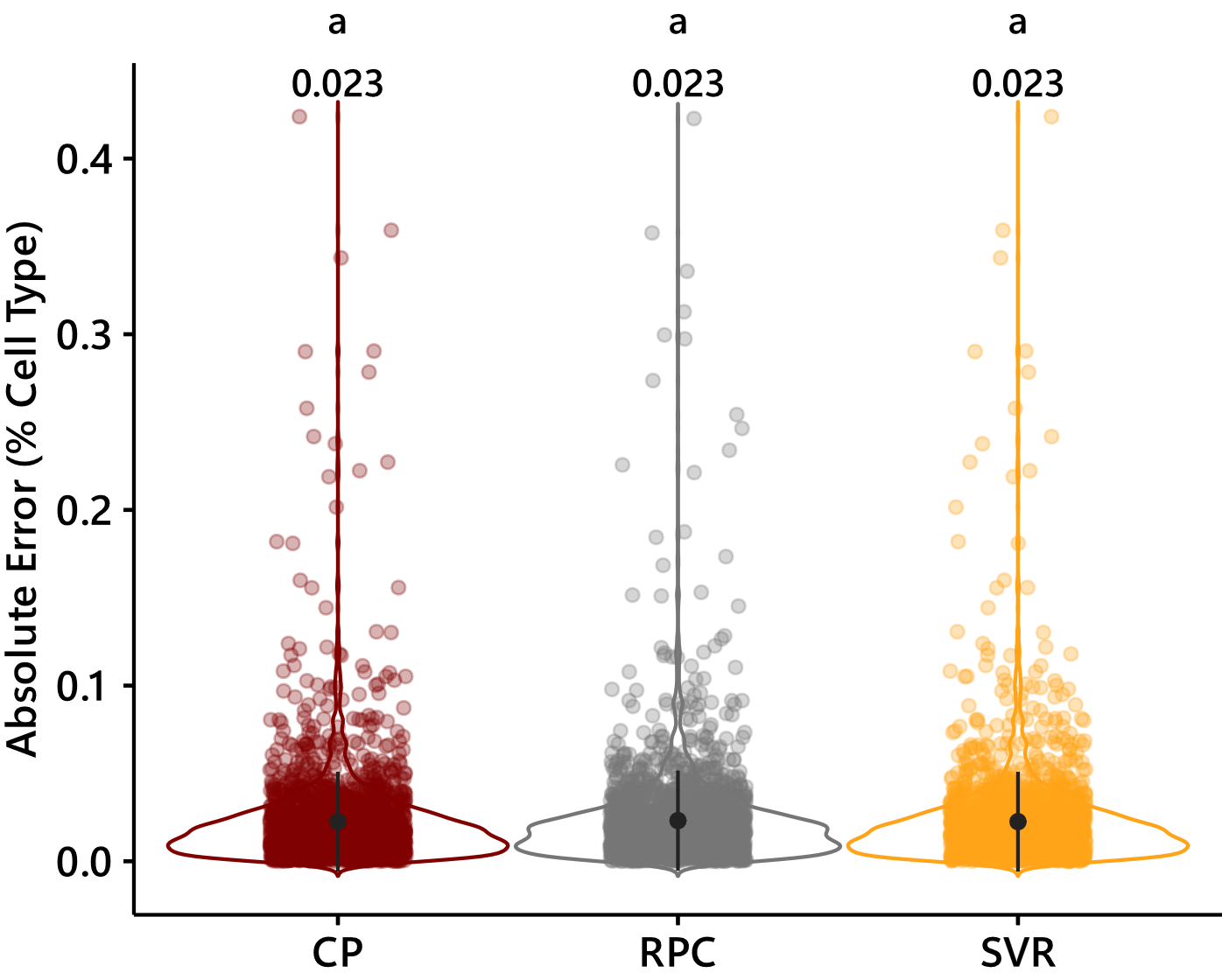

**Supplementary Figure 6.** Absolute prediction error of cell type proportions (predicted – true) across regression methods for proportion estimation. CP – constraint projection; RPC – robust partial correlation; SVR – support vector regression.
