## Supplementary Table 1 for "Advancing Pediatric and Longitudinal DNA Methylation Studies with CellsPickMe, an Integrated Blood Cell Deconvolution Method"

**Supplementary Table 1.** Pair-wise comparison of prediction error across normalization methods. Statistical significance was calculated with ANOVA with post-hoc Tukey HSD (Honestly Significant Difference) test. For each comparison, the normalization method that yielded significantly lowered AE is boldened and colored in navy blue.

| **group1** | **group2** | **statistic** | **df** | **p** | **p.adj** | **p.adj.signif** |
| --- | --- | --- | --- | --- | --- | --- |
| Noob | Funnorm | 2.1 | 25949 | 3.60E-02 | 3.60E-01 | **ns** |
| Noob | **Quantile** | 35.5 | 24830 | 3.26E-269 | 3.26E-268 | ******** |
| **Noob** | Quantile.B | -71.4 | 17923 | 0 | 0 | ******** |
| **Noob** | None | -81.9 | 18587 | 0 | 0 | ******** |
| Funnorm | **Quantile** | 33.5 | 24963 | 1.96E-240 | 1.96E-239 | ******** |
| **Funnorm** | Quantile.B | -72.8 | 17796 | 0 | 0 | ******** |
| **Funnorm** | None | -83.4 | 18446 | 0 | 0 | ******** |
| **Quantile** | Quantile.B | -92.7 | 16260 | 0 | 0 | ******** |
| **Quantile** | None | -105 | 16725 | 0 | 0 | ******** |
| **Quantile.B** | None | -4.79 | 25832 | 1.68E-06 | 1.68E-05 | ******** |
