## Supplementary Table 2 for "Advancing Pediatric and Longitudinal DNA Methylation Studies with CellsPickMe, an Integrated Blood Cell Deconvolution Method"

**Supplementary Table 2.** Percentage of predicted cell populations that fell within the clinical interval in Figure 4, out of the categories predicted. The normalization method that led to the highest percent of accurate predictions for each reference dataset is boldened.

| Median predicted proportions  (10th-90th Percentile) | Cord | UniBlood7 | IDOL | UniBlood13 | UniBlood19 | Extended | Average |
| --- | --- | --- | --- | --- | --- | --- | --- |
| Noob normalization (RGset) | 77.78% | 88.89% | **100.00%** | **63.16%** | **80.95%** | **83.33%** | 82.35% |
| Functional normalization (RGset) | 77.78% | 77.78% | **100.00%** | **63.16%** | 71.43% | 61.11% | 75.21% |
| Quantile normalization (RGset) | **100.00%** | **100.00%** | **100.00%** | 52.63% | 71.43% | 77.78% | 83.64% |
| Quantile normalization (Beta matrix) | 66.67% | 33.33% | 62.50% | 42.11% | 33.33% | 50.00% | 47.99% |
| No normalization | 55.56% | 55.56% | 37.50% | 42.11% | 33.33% | 38.89% | 43.83% |
| Average | 75.56% | 71.11% | 80.00% | 52.63% | 58.09% | 62.22% |  |
