## Supplementary Table 3 for "Advancing Pediatric and Longitudinal DNA Methylation Studies with CellsPickMe, an Integrated Blood Cell Deconvolution Method"

**Supplementary Table 3 –** Clinical interval of cell populations (10^th^ to 90^th^ percentile) in age 5-year healthy individuals of backgrounds across sexes and of various genetic ancestries. The cell counts were measured with flow cytometry or hematology analyzers.

|  | **Age-specific clinical intervals (10^th^-90^th^ Percentile)** | **Reference for clinical intervals 1 (DOI)** | **Age-specific clinical intervals 2 [Male and Female average] (10^th^-90^th^ Percentile)** | **Reference for clinical intervals 2 (DOI)** |
| --- | --- | --- | --- | --- |
| % B cells (% total lymphocytes) | 22.0  (12.9–29.2) | Tosato et al., 2015 (10.1002/cyto.a.22520) | 15.14  (10.21–21.77) | Ding et al., 2018 (10.1016/j.jaci.2018.04.022) |
| % T cells (% total lymphocytes) | 68.6  (59.7–77.6) |  | 67.60  (59.50–74.08) |  |
| % CD4+ T cells (% lymphocytes) | 38.0  (31.1–47.4) |  | 34.13  (26.17–41.07) |  |
| % CD8+ T cells (% lymphocytes) | 21.0  (16.0–26.9) |  | 26.13  (19.68–34.06) |  |
| CD4 / CD8 ratio | 1.77  (1.26–2.90) |  | 1.30  (0.87–2.05) |  |
| % NK cells (% total lymphocytes) | 8.0  (4.7–16.2) |  | 15.23  (7.83–22.24) |  |
| % Monocytes (% total WBC) | 3.8-10.5 | Bohn et al., 2023 (10.1111/ijlh.14068) |  |  |
| % Immature Granulocytes  (% total WBC) | 0.1-1.8 |  |  |  |
| % Granulocytes (% total WBC) | 14.4–86.1 |  |  |  |
| % Neutrophils (% total WBC) | 13.8-75 |  |  |  |
| % Basophils (% total WBC) | 0.1-0.6 |  |  |  |
| % Eosinophils (% total WBC) | 0.5-10.5 |  |  |  |
| % Nucleated RBC  (% nucleated cells) | 0-0 |  | 0.2-1.3 | Hwang et al., 2016 (10.1093/ajcp/aqv084) |
| % Transitional B cells (% B cells) | 6.3  (4.6-8.5) | Piątosa et al., 2010 (10.1002/cyto.b.20536) | 6.53  (2.58-12.3) | Ding et al., 2018 (10.1016/j.jaci.2018.04.022) |
| % Naïve B cells (% B cells) | 65.7  (47.3–77.0) |  | 63.43  (48.36–75.84) |  |
| % Memory B cells (% B cells) | 27.1  (18.6–46.7) |  | 13.55  (7.76–20.19) |  |
| % Naïve CD4+ T cells (% T cells) | 67  (46–99) | Schatorjé et al., 2012 (10.1111/j.1365-3083.2012.02671.x) | 62.53  (40.75–75.28) |  |
| % Memory CD4+ T cells (% T cells) | 20.1  (0.62–100) |  | 36.85  (23.85–59.86) |  |
| % Tregs | 8  (4–14) |  |  |  |
| % Naïve CD8+ T cells (% T cells) | 42  (16–100) |  | 61.65  (38.03–79.08) |  |
| % Memory CD8+ T cells (% T cells) | 52  (21–100) |  | 35.70  (14.84–71.29) |  |
